# Sequential sampling from memory underlies perceptual decisions unyoked from actions

**DOI:** 10.1101/2025.09.10.675475

**Authors:** Prayshita Sharma, Michael N. Shadlen, S. Shushruth

**Author notes:** Program in Computational Neuroscience, University of Chicago (Prayshita Sharma). **Contact:** (SS), (MNS).

## Abstract

Perceptual decision-making refers to the class of decisions in which sensory evidence is used to categorize percepts and guide actions. Conventionally, categorical decisions are thought to precede motor actions. However, recent studies in nonhuman primates challenge this assumption – when perceptual decisions were uncoupled from the actions they bear upon, animals postponed the decisions until relevant response options were revealed. To determine whether this postponement stems from cognitive limitations unique to nonhuman primates, we conducted a similar experiment with human subjects. Naive subjects viewed a random-dot motion (RDM) stimulus that was difficult to categorize. After a delay period following the RDM, two choice targets were presented and subjects decided which target lay closer to the perceived motion direction. Decision accuracy varied across subjects, reflecting individual differences in ability to integrate motion evidence. Notably, subjects with higher decision accuracy showed prolonged deliberation after choice-target presentation. Furthermore, the time they took to report their decisions depended on the strength of the motion evidence. This pattern of accuracy and decision reporting time could be accounted for by a bounded diffusion model in which subjects sequentially sample stored sensory information from memory during the target selection phase. When the RDM was challenging to categorize, the subsequent appearance of the targets provided a framework to interrogate stored evidence and render a decision. Our results reveal a strategic feature of working memory of retaining information based on its future utility. This observation opens new avenues for investigating how memory and decision-making interact.

## Introduction

Perceptual decisions intuitively feel like they precede and are separated from the actions they bear upon. For instance, a soccer goalkeeper might decide a shot is headed to their right (decide motion as “rightward”) and execute a dive in that direction (choose an action). But neurophysiological recordings in both animals and humans indicate that the accumulation of sensory evidence towards a decision transpires in the same neurons involved in action planning (Donner et al., 2009; Gallivan et al., 2018; Hanks & Summerfield, 2017; Selen et al., 2012; Shadlen & Newsome, 1996). This overlap suggests that perceptual decisions are embodied in terms of the motor actions that a percept affords (Cisek & Kalaska, 2010; Shadlen et al., 2008; Wispinski et al., 2020).

In behavioral tasks commonly used to study perceptual decision-making in the laboratory, the actions associated with decision outcomes are usually prespecified. It has been suggested that decisions are only embodied in terms of actions when such decision-to-action mapping is known in advance. In tasks in which decision-making is uncoupled from the actions they bear upon, “action-independent” decision signals have been reported in humans (Filimon et al., 2013; Gherman et al., 2024; O’Connell et al., 2012), monkeys (Bennur & Gold, 2011; Charlton & Goris, 2024; Freedman & Assad, 2006; Freedman et al., 2001; Genovesio et al., 2009; Gold & Shadlen, 2003; Goodwin et al., 2012; Horwitz et al., 2004; Quinn et al., 2021; Wang et al., 2019) and rodents (Pagan et al., 2025; Wu et al., 2020). However most of these studies still confined the subjects to choose between two possible actions for reporting their decision – primarily randomizing the mapping between the stimulus type and reporting action across trials. Thus, the degree of flexibility of the mapping between the decision and action uncovered by these studies is unclear. Further, in many of these studies, the stimulus that was being decided upon was unambiguous. Thus, it is difficult to ascertain the process by which the decision was formed.

A recent study in macaques (Shushruth et al., 2022) addressed these two issues. Monkeys were trained to associate two directions of stochastic random dot motion (RDM) stimuli with choice-targets of two colors. Importantly, the colored choice targets appeared following a delay after the RDM was extinguished at unpredictable locations. Thus, the monkeys had to learn to be highly flexible in mapping decisions to actions. And because of the stochastic nature of the RDM, the animals had to integrate motion evidence over time to arrive at a decision – and this deliberation would serve as a signature of the decision making process. Surprisingly, this study found that monkeys, instead of integrating evidence towards a decision when the RDM was shown, postponed it until after the choice targets were shown. Further, activity of neurons in the sensorimotor association area LIP represented decision formation during this action selection epoch. These results suggested that the monkeys were storing sensory information during RDM viewing and retrieving it from memory to evaluate it in the context of available responses.

The postponement of evidence accumulation could arise from an inability of monkeys to effectively categorize the stimulus, possibly due to their limited conceptual understanding of direction. Alternatively, when sensory evidence is stochastic and thus the percept is uncertain, delaying commitment until the decision can be instantiated in the appropriate action framework may be advantageous (Drugowitsch et al., 2016). We hypothesized that when confronted with stimuli that are difficult to categorize, humans also resort to a strategy similar to that used by monkeys. To test this, we designed a decision-making task in which noisy motion evidence makes direction difficult to categorize. Human subjects estimated the net direction of an RDM which varied over 360° across trials. After a delay period following the RDM, two choice targets were shown. Subjects indicated their decision by choosing the target closest to their perceived motion direction.

The decision accuracy varied across the six subjects, reflecting differences in their ability to integrate motion evidence. Notably, subjects with higher decision accuracy showed prolonged deliberation after choice-target presentation, and the time they took to report their decisions depended on the strength of the motion evidence. We found that this pattern of accuracy and decision reporting time can be explained by a model in which subjects sequentially sample stored information from memory while choosing the targets (Ratcliff & Rouder, 1998; Shadlen & Kiani, 2013; Shadlen et al., 2006; Shushruth et al., 2022). When the stimulus was hard to categorize, the appearance of the targets provided a framework to interrogate the stored evidence and render a decision. Our results show that working memory retains sensory details strategically, based on their anticipated future use.

## Materials and Methods

### Subjects

Six adult subjects (three males and three females) participated in the experiment after providing informed written consent. The experiments were approved by the Institutional Review Board of Columbia University (IRB #AAAL0658). All the subjects were naive to the hypothesis of the experiment and five out of the six subjects had no previous experience with random dot direction discrimination tasks. Each subject performed the task over one or two sessions with a maximum duration of 3 hours each, including breaks. Not included is data from two subjects (S1 and S2), who performed pilot versions of the task and one subject (S8) who withdrew consent.

#### Justification of the sample size

The sample size reflects the within-subject design of the experiment. All model fitting and key behavioral analyses were performed independently for each subject, allowing decision-making strategies to be characterized without relying on population averaging. Importantly, the core behavioral phenomenon was observed in all six subjects, with inter-individual variability reflected primarily in the strength of the effect and fitted model parameters. This combination of dense within-subject sampling and relatively small subject numbers is consistent with recent perceptual decision-making studies that fit bounded evidence-accumulation models at the individual-subject level (Kira et al., 2025; Smith & Corbett, 2019; Stine et al., 2020).

### Apparatus

Subjects were seated in a comfortable chair in a semi-dark room, with their chin and forehead stabilized on a tower mounted on a table. They viewed visual stimuli on a ViewPixx monitor (60 Hz refresh rate, 68 cm viewing distance). Their eye movements were recorded at 1 kHz with a non-invasive infrared eye-tracking device (Eyelink 1000, SR Research Ltd., Ontario, Canada). The temporal precision of the visual stimuli was controlled by PsychToolBox (www.psychtoolbox.org) run on a Macintosh computer.

### Behavioral tasks

Subjects made decisions about the net direction of stochastic random-dot motion (RDM) stimuli across multiple trials (Figure 1). Each trial started after the subject fixated on a circular fixation point (FP) on the screen. After a variable delay (400 − 800 ms, drawn from a truncated exponential distribution), the RDM stimulus was displayed in a circular aperture of radius 5^◦^ visual angle around the FP. The RDM stimulus was based on previous studies (Roitman & Shadlen, 2002), but modified to increase the difficulty of direction estimation. On each trial, a motion direction *θ* ∈ {0, 10, 20, …, 350} and a motion strength *C* ∈ {0, 0.04, 0.08, 0.16, 0.32, 0.64} were randomly chosen. The first three frames of the stimulus consisted of white dots randomly plotted at a density of 16.7 dots· deg^−2^ · s^−1^. From the fourth frame, a fraction of dots *C* from three frames before were randomly chosen to be ‘signal dots’. Each signal dot was displaced in a direction randomly sampled from a von Mises distribution 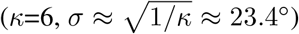 centered at *θ*. The remaining dots were replotted at random locations. The stimulus was presented for a variable duration drawn from a truncated exponential distribution (range 250–800 ms, mean 350 ms). 400 ms after the cessation of the stimulus, the FP was extinguished and two circular targets were presented at an eccentricity of 6^◦^ visual angle from the center, one in the direction *θ* (‘correct’ target) and another at *θ* ± 40^◦^ (‘incorrect’ target). Subjects reported their decision by making an eye movement to the target closest to their perceived direction of motion. We refer to the time between target presentation and decision report as go-reaction time (*go-RT*). A subset of three subjects also performed the task with a 1200 ms interval between RDM cessation and target onset. To ensure that subjects were motivated to engage with this challenging task, we offered them a performance-based monetary bonus.

**Figure 1.**
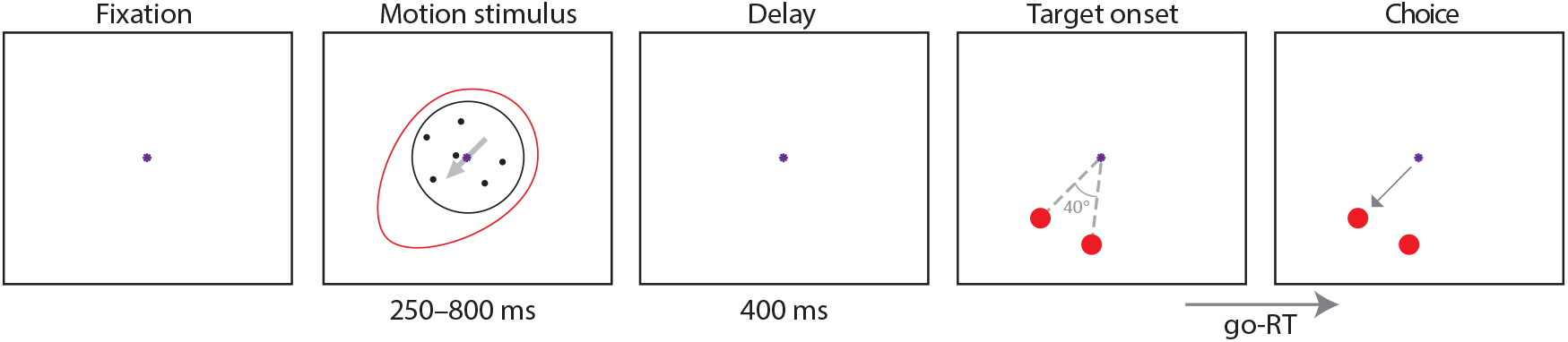
Direction-decision task. After subjects acquire fixation, a random dot motion stimulus is presented. The direction of motion of the coherent dots is drawn from a von Mises distribution (denoted by the red oval - not visible to the subject). Across trials, the mean of the distribution varies randomly between 0°-350° (in 10° increments). This is followed by a short delay period. Two targets are then presented equidistant from the fixation point. One target is in the direction of the mean of the von Mises and the other, 40° away. The subjects report their choice by an eye movement to one of the two targets.

### Analysis

The accuracy of the subjects’ decisions (Figure 3) were fit with a logistic model of the following form:

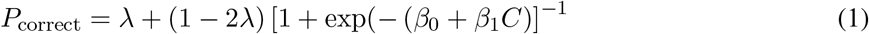

where *λ* (lapse rate, i.e. errors at highest coherence), *β*_0_ (bias) and *β*_1_ (sensitivity) are fit parameters.

**Figure 2.**
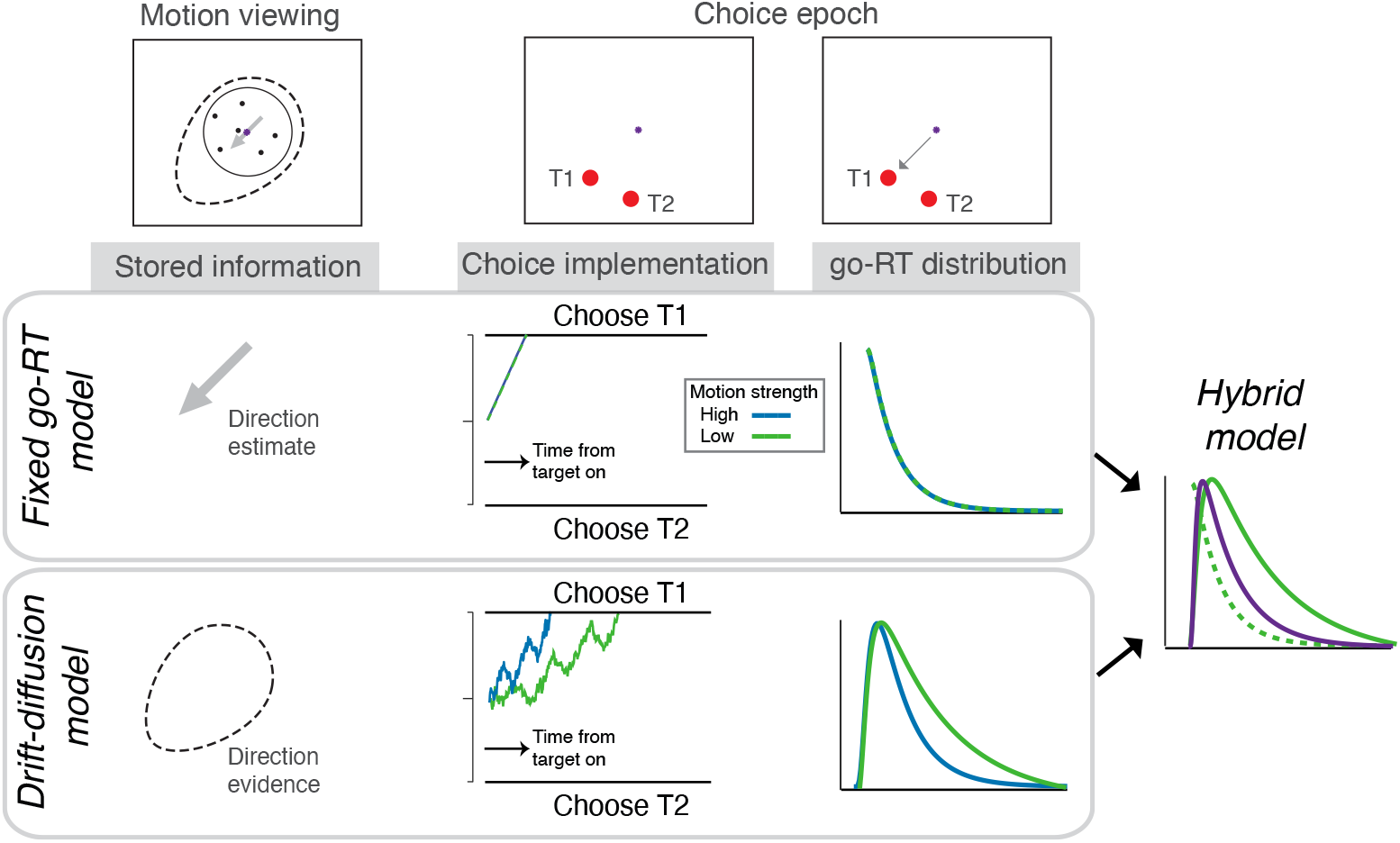
Schematic of decision models. In the *fixed go-RT model*, the subjects decide on the direction during motion viewing and store a scalar estimate in memory. When the choice targets appear, this stored estimate is directly mapped onto the nearest target, yielding fast go-RTs that are largely invariant to motion coherence. In the *drift-diffusion model*, subjects store a representation of the sensory evidence itself during motion viewing. After the onset of choice targets, the evidence in memory is sequentially sampled to render a choice. This results in slower go-RTs that systematically vary with coherence. In the *hybrid model*, subjects are assumed to flexibly deploy these two strategies across trials, favoring rapid choices when motion evidence is strong, and memory-based evidence accumulation when direction estimation is uncertain.

**Figure 3.**
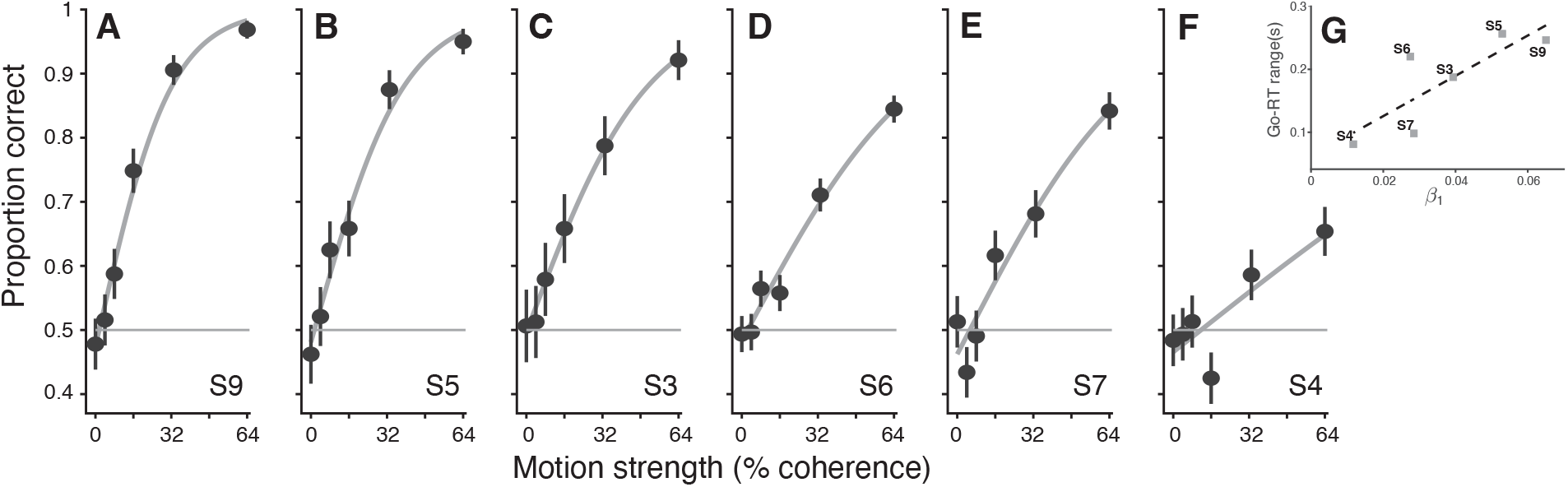
Decision Accuracy. **(A-F)** The proportion of correct choices is plotted as a function of motion coherence for individual subjects. Solid lines are fits to a logistic model (Equation 1). The subjects are in the order of decreasing sensitivity (*β*_1_ from logistic fits). **(G)** Correlation between *β*_1_ and range of go-RT across the six subjects.

To assess the correlation between *β*_1_ and go-RT across subjects, we computed the metric *go-RT range* as the difference in mean go-RT between the lowest and highest coherence condition for each subject.

### Model fitting

To investigate the relation between subject’s choices and go-RTs, we fit the distribution of go-RTs with a variety of computational models (Figure 2). Each model provided predicted go-RT probability densities for each coherence level. The models were fit by maximizing the log likelihood of the observed go-RT distributions given each model’s predicted distribution. For models that also provided predictions of choice distributions, fitting was performed by jointly maximizing the log likelihood of both choice and go-RT distributions under the model. 95% confidence intervals for the model parameters were computed by bootstrap resampling the observed trials and refitting the model.

### Drift-diffusion model

We first fit choice accuracy and go-RTs of individual subjects with bounded drift-diffusion (DtB) models (Shadlen et al., 2006). Traditionally, for direction discrimination tasks, perceptual samples of evidence are considered to be accumulated during RDM viewing. In our adaptation of the model, the samples of evidence are instead presumed to be stored in memory during the RDM viewing epoch. When the target locations are revealed, these samples in memory are integrated to decide which target to choose. The evidence from memory at each time step is assumed to arise from a normal distribution with variance Δ*t* and mean *κ* ∗ *C* ∗ Δ*t*, where *C* is the motion coherence, and *κ* is a scaling parameter. The samples of instantaneous evidence are assumed to be independent and this evidence is accumulated as a decision variable (DV), until the integrated evidence reaches an upper or lower bound (±*B*) which determines the subject’s choice. The go-RT would be the time taken to reach the bound, together with a non-decision time (*T*_*nd*_) to account for sensory and motor delays.

We first considered two versions of the decision bound: a fixed bound *B*, and a bound that collapses over time as follows (Drugowitsch et al., 2012):

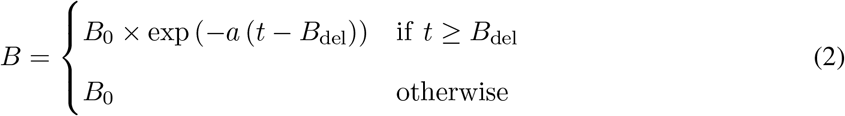

where *B*_0_ is the initial bound height, *a* is the rate of collapse and *B*_del_ is the delay to onset of collapse. Formal model comparison using BIC favored the fixed bound model for all subjects, and subsequent models and analyses therefore use this simpler bound formulation. Stimulus duration was not explicitly included as a parameter in this model of post go-cue decision process.

We fit the drift-diffusion model to the choice and go-RT data using a maximum likelihood procedure. The go-RT and decision accuracy for a given set of parameters were computed from numerical solutions to Fokker-Planck equations (Chang & Cooper, 1970; Kiani & Shadlen, 2009).

### Fixed go-RT model

We considered an alternative model in which subjects reach a decision during RDM viewing or delay epoch, and report their decision after target onset (after sensory and motor delays). This model assumes that the mean go-RTs are invariant to the strength of motion (Figure 2).

The trial-to-trial variability of go-RTs was modeled as an exponential distribution (Luce, 1991):

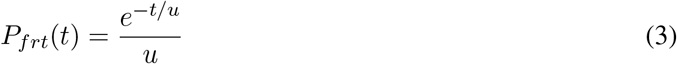

where *t* represents time following target onset and a sensorimotor latency, and *u* is the mean go-RT. Note that the fixed go-RT model does not directly predict choice distributions, as it assumes choices and go-RTs are independent.

### Hybrid Model

We formulated a hybrid model to capture the possibility that subjects switch between decision-making during RDM and delayed integration, based on the reliability of motion evidence on each trial. The contributions of the diffusion model and fixed go-RT model to the go-RT distribution are captured by a coherence dependent scaling factor, *α*, that is fit to each subject. The go-RT probability distribution in the model for coherence *C* is:

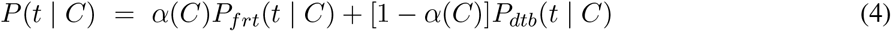

where *P*_*frt*_(*t* | *C*) and *P*_*dtb*_(*t* | *C*) are, respectively, go-RT probabilities from the fixed go-RT model and the bounded drift diffusion model, and *α*(*C*) ∈ [0, 1] is the fractional contribution of the fixed go-RT model.

To model the increasing contribution of the fixed go-RT model as a function of motion strength, we parameterized the scaling factor as a linear function of coherence: *α*(*C*) = (1 − *s*) + *s C, s* ∈ [0, 1]. Here, the parameter *s* governs how strongly the fast-strategy is favored as coherence increases. At *s* = 0, *α*(*C*) = 1 and the model reduces to a pure fixed go-RT strategy; at *s <* 1, the contribution of the fixed go-RT model increases linearly with coherence.

This hybrid formulation provides a flexible yet parsimonious account of how decision strategy can vary with stimulus strength, with the slope parameter *s* capturing the shift from decisions during RDM viewing to integration from memory at different levels of motion strength.

### Comparison between models

We used Bayesian Information Criterion (BIC) to adjudicate between the drift-diffusion, fixed go-RT and hybrid models. Since the fixed go-RT model can only explain go-RTs but not choice, the comparison between drift-diffusion and fixed go-RT models were made solely from fitting go-RTs. For comparing the drift-diffusion to hybrid models, we used the complete fits to both choice and go-RT distributions.

To assess whether the candidate models could be distinguished given the number and distribution of trials in our experiment, we performed model recovery analyses. For each candidate model, we simulated 200 datasets using the subjects’ best-fitting model parameters. Each simulated dataset was then fit with all three candidate models using the same fitting and model-comparison procedures applied to the empirical data. The recovered model was defined as the model with the lowest BIC. Recovery performance is shown as a confusion matrix in Table 1.

**Table 1:** Model recovery. Rows indicate the model used to generate synthetic datasets and columns, the model selected by BIC. Values are proportions of simulated datasets.

| Generating model | Recovered model |  |  |
| --- | --- | --- | --- |
|  | Drift-diffusion | Fixed go-RT | Hybrid |
| Drift-diffusion | 0.78 | 0.22 | 0.01 |
| Fixed go-RT | 0.00 | 1.00 | 0.00 |
| Hybrid | 0.22 | 0.12 | 0.67 |

The candidate models were distinguishable, although not perfectly so. The fixed go-RT model and driftdiffusion models were recovered on 100% and 78% of their respective simulations. While the hybrid model was correctly recovered in 67% of simulations, data generated by the drift-diffusion and fixed go-RT models were almost never classified as hybrid (1% and 0% of simulations, respectively). Thus, model recovery was conservative with respect to identifying a hybrid process.

### Effects of prolonging delay epoch

In a second experiment conducted on a subset of three subjects in separate sessions, on 50% of the trials, the time between RDM cessation and target onset was extended from 400 ms to 1200 ms. To test whether the duration of the delay significantly affected the slope of the logistic fits, we included an interaction term in the logistic regression model in Equation 1 as follows:

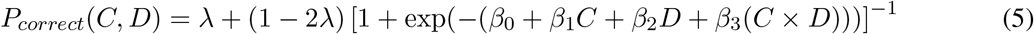

where *β*_2_ represents the main effect of delay duration *D* (coded as 0 for 400 ms and 1 for 1200 ms), and *β*_3_ captures the interaction between coherence and delay duration. The significance of the interaction effect was tested by evaluating whether *β*_3_ was significantly different from zero. Models with and without the interaction term were also compared using a Bayes factor.

## Results

Six naive human subjects performed a direction discrimination task (Figure 1). They were first presented a random-dot motion (RDM) stimulus whose dominant direction varied between 0^◦^ − 350^◦^. After a 400 ms delay following stimulus offset, two targets appeared equidistant from fixation – one aligned with the motion direction and the other offset by ±40^◦^. subjects were instructed to choose the target aligned with the motion direction. The difficulty of direction discrimination varied across trials as noise was introduced to both motion direction and fraction of coherently moving dots (see Materials and Methods).

### Subjects exhibit deliberation during action selection

All six subjects found the task challenging. Unlike standard RDM tasks where subjects rarely make errors at high coherence (*lapses*), all subjects in this task showed lapses at 64% coherence trials (Figure 3). Since, to correctly discern the net direction of motion in this task, one needs to integrate motion information over time (Kiani et al., 2008; Stine et al., 2020), the presence of lapses underscores the demand the task placed on effective integration of motion evidence. Subjects varied notably in their sensitivity to motion coherence – those with fewer lapses exhibited higher sensitivity to motion strength. This suggests that subjects varied in their ability to integrate motion evidence over time.

The variability across subjects in the ability to integrate motion information could have arisen in two ways. One possibility is variability in perceptual sensitivity to motion stimuli during the viewing phase, a hypothesis partially supported by logistic regression coefficients. Intriguingly, recent studies in monkeys (Shushruth et al., 2022; Wang et al., 2019) suggest another possibility – when motion evidence is weak, the subjects may store the motion information in memory and later, once the targets are presented, use the location of the targets as spatial reference to integrate the stored motion information to decide which target to choose.

In monkeys, a signature of evidence integration from memory was prolongation of the time between target onset and decision report (go-reaction time, go-RT, Figure 1). Consistent with this, the accuracy of our subjects’ decisions (slope of logistic fit in Figure 3, Equation 1) was correlated with the range of go-RT between the easiest and the hardest decisions (*r* = 0.82, *p* = 0.047, Pearson’s correlation, Figure 3G). More importantly, subjects with higher accuracy also showed a systematic dependence of go-RT on motion coherence (Figure 4), mirroring patterns observed in monkeys (Shushruth et al., 2022). This coherence-dependent increase in go-RT is suggestive of deliberation about stored motion evidence after the targets appear.

**Figure 4.**
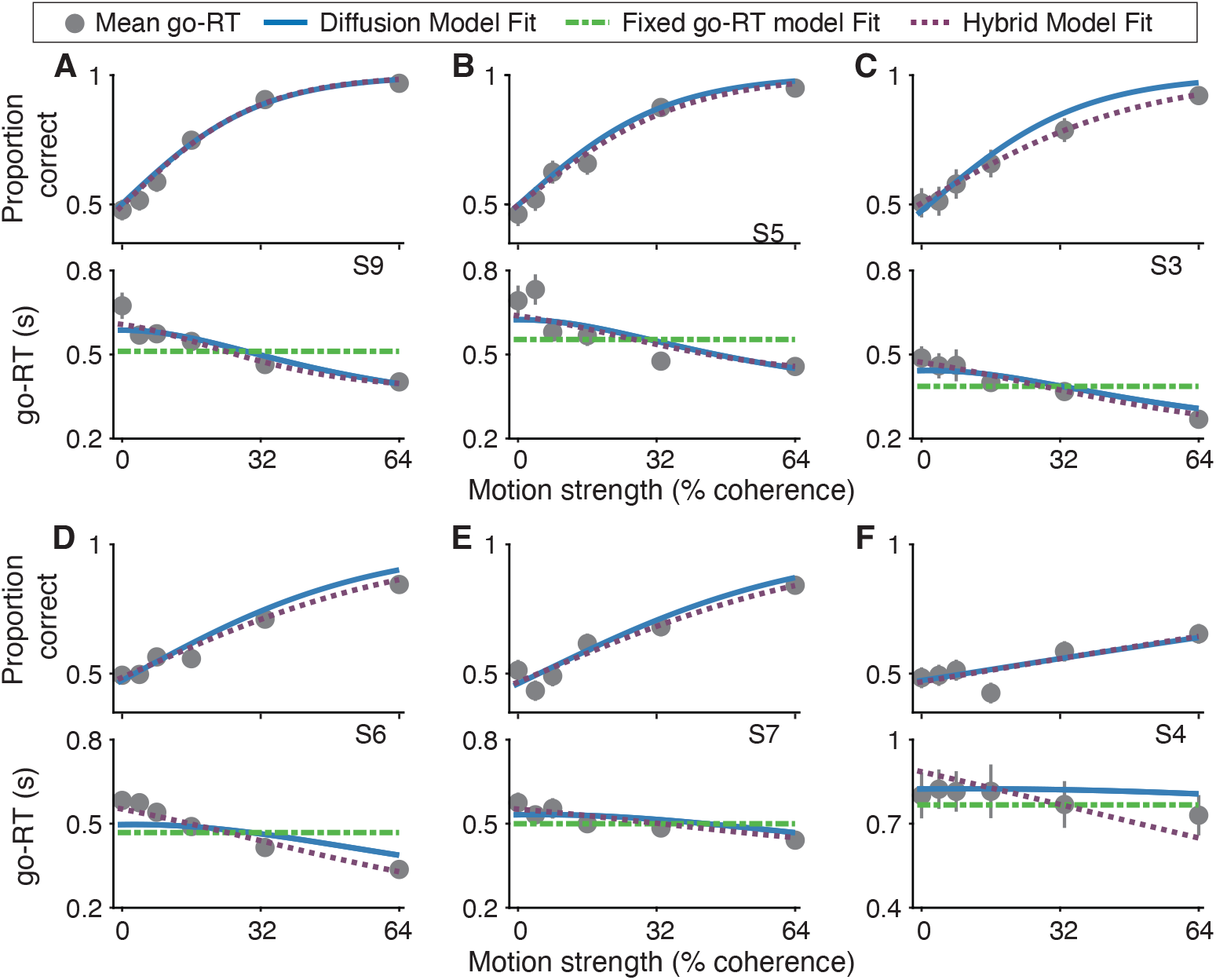
Model fits for decision accuracy and go-RT. For each subject, the proportion of correct choices (*top*) and go-RTs (*bottom*) are plotted as a function of motion strength. Lines are fits to various models indicated in the legend. Error bars indicate SEM. Subjects are ordered as in Figure 3.

### Drift-diffusion models account for deliberation during action selection

We tested whether prolonged go-RTs enhanced decision accuracy by modeling subjects’ choice and go-RT data using a diffusion-to-bound (DtB) framework. In DtB models, sensory evidence is accumulated as a decision variable and a choice results when the decision variable reaches one of two boundaries. Typically, reaction times in a standard RDM task correspond directly to evidence accumulation duration, providing a powerful mechanistic link between choice accuracy and RT. In our task, if subjects integrated evidence after targets appeared, a DtB model would similarly link choices and prolonged go-RTs.

Indeed, DtB models provided robust fits to individual subjects’ choice data and go-RT distributions (Figure 4, blue lines). The model’s performance was especially strong for subjects showing pronounced go-RT dependence on coherence, who also exhibited higher decision accuracy. We first compared these fits to an alternative model that assumes subjects form an abstract decision during motion viewing and quickly convert that decision into a choice when the targets are shown (*fixed go-RT model*, Equation 3). In this model, go-RT is independent of motion coherence, and go-RT variability is attributed to motor and other non-decision processes. Because the fixed go-RT model does not account for the dependence of choice on coherence, model comparisons were restricted to the go-RT distributions. Using Bayesian Information Criterion (BIC), these comparisons strongly favored the DtB model for most subjects, except subject S4, who exhibited the lowest decision accuracy (Figure 5A). This supports the idea that go-RT structure reflects decision deliberation during action selection.

**Figure 5.**
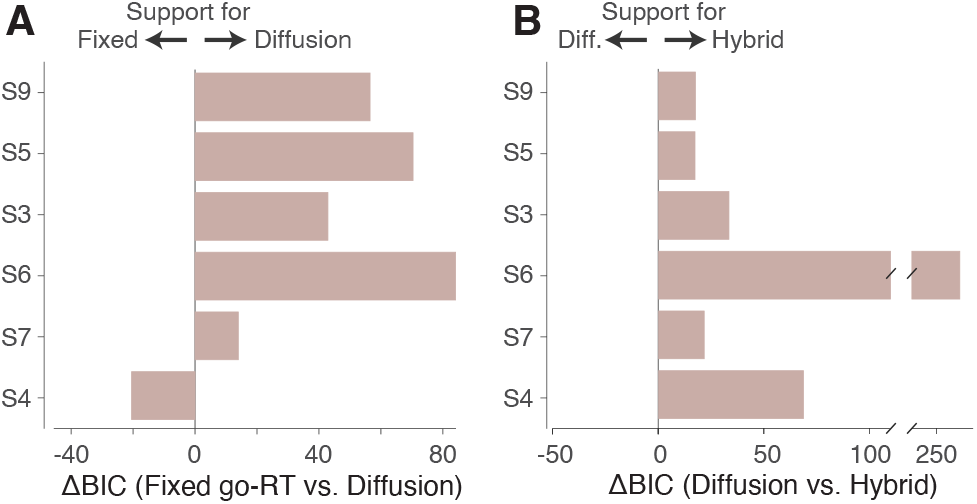
Model comparisons. Δ*BIC* for comparing drift-diffusion model (DtB) to either **(A)** the fixed go-RT model or **(B)** the hybrid model. Each bar represents individual subjects. Subjects are ordered as in Figure 3.

To explore individual differences in strategy further, we analyzed model fits for each subject across different coherence conditions (Figure 6). For most subjects, the DtB model outperformed the fixed go-RT model primarily at lower coherence conditions. This led us to consider a hybrid, two strategy model. In this model, subjects make an abstract decision when motion evidence is strong but resort to evidence integration from memory of the stimulus when motion evidence is ambiguous. The relative contributions of DtB vs. fixed go-RT model to the go-RT distributions were fit per individual subject with a scaling factor (*α*, Equation 4). Because both the hybrid and DtB models make predictions about both choice and go-RT, this second comparison was performed on the joint fits to both data types. This hybrid model (purple dotted line in Figure 4 and Figure 6) provided superior fits over the diffusion model for all subjects (Figure 5B).

**Figure 6.**
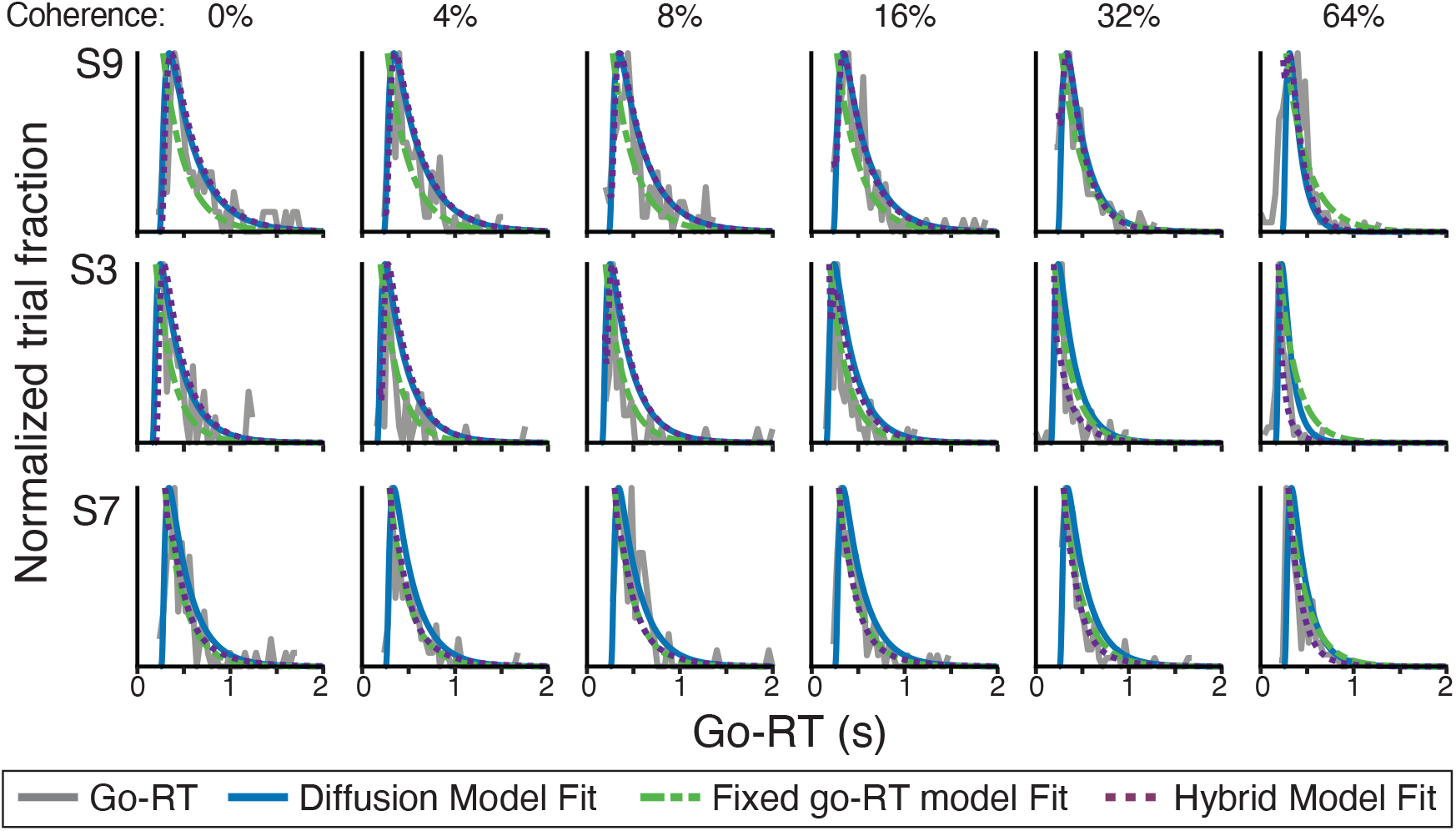
Distributions of go-RTs and model fits. For three example subjects, the distribution of go-RTs for correct choices are plotted at different coherences (gray lines). Superimposed are the fits to the three models indicated in the legend.

These results demonstrate a strategic use of memory during decision-making, with subjects storing either an abstract estimate of motion direction (when evidence is strong) or samples of motion evidence itself (when ambiguous), enabling flexible deliberation once targets are revealed. While our psychophysical data establish that subjects can sequentially sample motion information from memory, they do not adjudicate the precise form of the stored representation or the decision rule applied after sampling (such as graded accumulation or extrema detection) (Stine et al., 2020). The fitted *s* values of individual subjects provided insight into how different individuals weighted fast vs. deliberative decisions as a function of coherence (Table 2), underscoring individual differences in memory utilization and decision-making approaches.

**Table 2:** Model fit parameters (with 95% confidence intervals) for individual subjects.

|  | DtB |  |  | Fixed go-RT |  |
| --- | --- | --- | --- | --- | --- |
| | $\kappa$ ($coh^{-1} s^{-1}$) | $B$ | $T_{nd}$ (s) | $u$ (s) | $T_{nd}$ (s) |
| S9 | 5.30 [4.52, 6.25] | 0.57 [0.54, 0.59] | 0.23 [0.21, 0.24] | 0.24 [0.21, 0.27] | 0.28 [0.26, 0.29] |
| S5 | 5.08 [4.15, 5.94] | 0.55 [0.52, 0.57] | 0.28 [0.27, 0.29] | 0.23 [0.20, 0.25] | 0.33 [0.32, 0.34] |
| S3 | 5.32 [3.87, 6.80] | 0.50 [0.46, 0.54] | 0.15 [0.14, 0.17] | 0.19 [0.16, 0.22] | 0.20 [0.19, 0.21] |
| S6 | 2.91 [2.52, 3.39] | 0.59 [0.58, 0.61] | 0.10 [0.10, 0.11] | 0.31 [0.28, 0.33] | 0.17 [0.16, 0.17] |
| S7 | 3.06 [2.44, 3.72] | 0.50 [0.47, 0.52] | 0.25 [0.24, 0.26] | 0.21 [0.18, 0.23] | 0.30 [0.29, 0.30] |
| S4 | 0.65 [0.35, 1.11] | 0.79 [0.75, 0.81] | 0.14 [0.13, 0.16] | 0.50 [0.44, 0.50] | 0.27 [0.25, 0.29] |

|  | Hybrid |  |  |  |
| --- | --- | --- | --- | --- |
| | $\kappa$ ($coh^{-1} s^{-1}$) | $B$ | $T_{nd}$ (s) | $s$ |
| S9 | 5.38 [3.90, 6.61] | 0.57 [0.54, 0.61] | 0.24 [0.23, 0.35] | 1.00 [0.59, 1.00] |
| S5 | 4.16 [3.31, 5.36] | 0.61 [0.56, 0.66] | 0.35 [0.33, 0.38] | 0.63 [0.48, 0.83] |
| S3 | 3.70 [2.29, 5.01] | 0.49 [0.47, 0.61] | 0.19 [0.19, 0.27] | 1.00 [0.47, 1.00] |
| S6 | 2.14 [1.81, 2.78] | 0.67 [0.64, 0.69] | 0.24 [0.23, 0.24] | 0.65 [0.60, 0.68] |
| S7 | 2.26 [1.69, 3.95] | 0.59 [0.36, 0.65] | 0.30 [0.27, 0.31] | 0.50 [0.34, 0.94] |
| S4 | 0.53 [0.28, 1.27] | 1.04 [0.32, 1.11] | 0.30 [0.27, 0.34] | 0.40 [0.27, 0.51] |

### Motion information is held in working memory

The RDM stimulus is noisy and stochastic. How can subjects store such sensory information over hundreds of milliseconds? One possibility is that this information is held in iconic memory, which is considered a property of sensory systems (Sperling, 1960). While iconic memory is high capacity, it is also labile and decays rapidly within a second (Graziano & Sigman, 2008). For intermediate-term storage, some aspects of iconic memory are thought to be transferred to working memory, but this storage is known to be capacity limited.

To test whether the stimulus information driving decisions is held in working or iconic memory, we re-tested the three most proficient subjects in a modified task version, where the wait duration on each trial could be either 400 or 1200 ms. We focused on these subjects because the comparison is most informative in participants who clearly performed the task by integrating motion evidence, particularly at lower coherence.

In all three subjects, we found no evidence for a reduction in decision sensitivity at the longer 1200 ms wait duration (Figure 7, ℋ_0_: *β*_3_ = 0 in Equation 5; Table 3). Thus, in subjects who clearly performed the task by integrating motion information, the relevant representation remained usable over a delay well beyond the timescale usually associated with iconic memory, consistent with the involvement of working memory.

**Table 3:** Logistic regression interaction coefficient (*β*_3_) for the coherence × delay interaction, 95% confidence intervals, and Bayes Factors (BF_01_)

| | $\beta_3$ | $p$ | 95% CI | BF<sub>01</sub> |
| --- | --- | --- | --- | --- |
| S9 | -2.20 | 0.28 | [-6.26, 1.87] | 8.64 |
| S5 | -1.37 | 0.65 | [-7.37, 4.63] | 13.74 |
| S3 | 1.17 | 0.63 | [-3.67, 6.00] | 11.84 |

**Figure 7.**
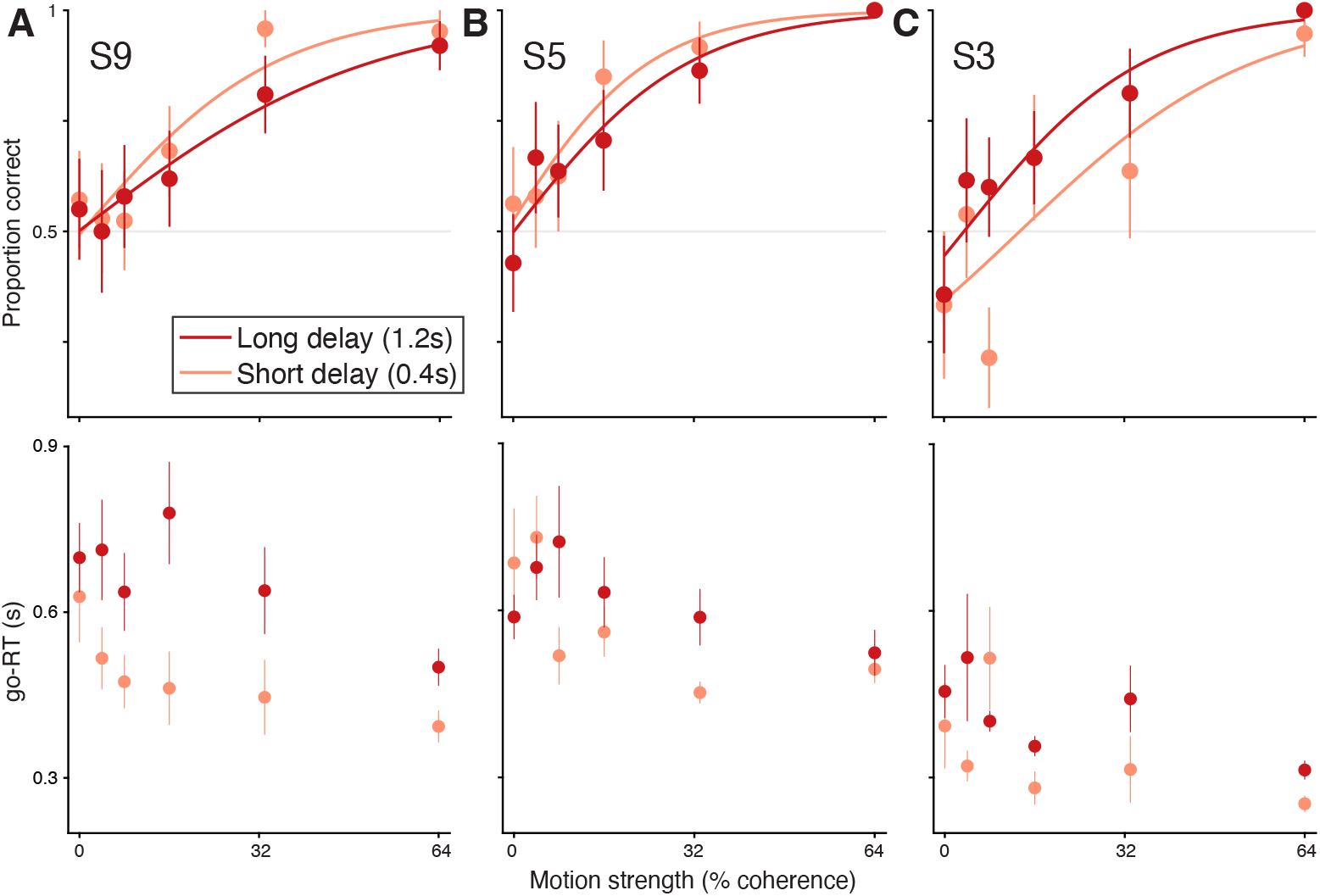
Choice and go-RT distributions at different memory delays. Proportion of correct choices (top) and go-RTs (bottom) are plotted as a function of motion coherence for three subjects tested with two different delays between motion offset and target onset. Solid lines indicate logistic fits to the choice data (Equation 1).

Although choice accuracy was similar across the two memory delays in these subjects, the pattern of go-RTs was more complex. Go-RTs continued to vary with motion coherence for both wait durations, consistent with the use of stored motion information during action selection. In addition, longer wait durations were associated with overall increases in go-RT. Although the number of trials in this experiment precluded a deeper analysis of the relation between prolonged go-RTs and preserved choice accuracy, the overall pattern suggests that increasing the memory delay altered the temporal dynamics of decision formation without measurably reducing performance.

## Discussion

The study of perceptual decision making has offered a window into cognition by revealing the processes through which sensory information is deliberated upon to guide motor actions (Kiani et al., 2014). A salient result from this line of research is that the accumulation of sensory evidence for decisions is instantiated by neurons associated with motor planning (de Lafuente et al., 2015; Ding & Gold, 2010, 2012; Roitman & Shadlen, 2002). This observation has led to the proposal that perceptual decision-making is embodied as a choice between potential actions (Cisek, 2007; Shadlen et al., 2008). However, this framework typically applies when actions for reporting decisions are specified in advance. The current study demonstrates that decision-making can remain embodied even when response actions are specified post-stimulus. Specifically, when perceptual information is ambiguous and requires temporal integration, subjects can maintain the information in working memory and strategically delay evidence accumulation until the appropriate response actions are presented. We discuss the relevance of our results to the concept of embodied cognition and to our understanding of working memory.

### Implications for embodied cognition

The concept of embodied perceptual decision-making has been widely debated (Cisek & Pastor-Bernier, 2014; Wispinski et al., 2020). Decision related signals have been reported in humans and animals making perceptual decisions without explicit knowledge of decision-action mapping (Bennur & Gold, 2011; Filimon et al., 2013; Freedman & Assad, 2006; Genovesio et al., 2009; Gherman et al., 2024; Goodwin et al., 2012; Horwitz et al., 2004; O’Connell et al., 2012; Pagan et al., 2025; Quinn et al., 2021; Wu et al., 2020). However most of these studies were structured to randomize between two available decision-to-action mappings. This leaves open the possibility that subjects made provisional motor decisions and adjusted them when the stimulus response associations were revealed. For instance, in spatial orienting paradigms, reorienting a planned saccade can occur rapidly, on the order of a few tens of milliseconds (Rizzolatti et al., 1987; Walker et al., 1995). Our task substantially reduces this possibility by making it difficult to prepare the final motor response during stimulus viewing.

Prior studies have shown that, in tasks similar to the one in the current study (Shushruth et al., 2022; Wang et al., 2019), nonhuman primates postpone decision-making until the decision-action mapping is revealed. Our results suggest that humans also often postpone decision formation until action selection, as evidenced by prolonged go-reaction times that scale with motion strength. However, postponed deliberation should not be expected in all delayed-report tasks. We suggest that it is most likely when two conditions are met. First, the stimulus is sufficiently uncertain that decision-makers who prioritize accuracy over speed benefit from delaying commitment. Second, the decision-to-action mapping cannot be reduced during stimulus viewing to a small, pre-specifiable set of action plans across which subjects can rapidly reorient. When these conditions are not met, as in tasks with more limited stimulus-response ambiguity (Coallier & Kalaska, 2014; Twomey et al., 2016), subjects may complete the decision before target onset. When they are met, as in the present study and in Bang and Fleming (2018), post-target deliberation becomes more likely.

Our results reinforce the view that, despite humans’ capacity for abstract cognition, the organization of perceptual knowledge fundamentally serves potential actions (Clark, 1997; Gibson, 1979; Merleau-Ponty, 1962). Thus, when faced with perceptual stimuli that resist easy categorization, humans appear to default to an embodied strategy, using action-oriented frameworks to guide decision formation. Such a strategy can help mitigate some of the known suboptimalities humans exhibit during decision-making in uncertain environments. (Drugowitsch et al., 2016)

### Working memory representations of sensory information

There is a rich history of psychophysical work characterizing drift-diffusion models in the context of working memory retrieval. (Forstmann et al., 2016; Ratcliff & Rouder, 1998; Ratcliff et al., 2016) In this literature, drift-diffusion processes are typically invoked to describe decisions about the contents of memory itself, including retrieval of single items (Smith, 2016) and multi-item cued recall (Schneegans & Bays, 2016). Our results extend this framework by demonstrating that working memory can also serve as a substrate for storing sensory evidence for deferred perceptual decision-making.

Given that sensory evidence from RDM stimulus is noisy and stochastic, it would seem that the memory requirements that can support protracted integration of such evidence are daunting. This information could reside in iconic short term memory which is a high capacity memory buffer (Sperling, 1960). It has been demonstrated that such memory can be formed strategically in anticipation of operations that may be required (Gegenfurtner & Sperling, 1993; Sperling & Weichselgartner, 1995). However, iconic memory is highly labile and decays rapidly (Graziano & Sigman, 2008). Indeed, our modified experiment with extended delay duration showed that stimulus information is held for longer than a second, suggesting the involvement of the working memory system.

Working memory is classically thought to be of limited capacity (Luck & Vogel, 1997), although more recent work has revealed its more dynamic and adaptive properties (Buschman & Miller, 2022; Ma et al., 2014). In RDM tasks, it is possible to strategically store limited samples of evidence to support good accuracy (Kang et al., 2021; Shushruth et al., 2022; Stine et al., 2020), making working memory a feasible site of storage. An emerging perspective is to consider the purpose of working memory as not mere retention of information, but to make information readily available for cognitive computations (van Ede et al., 2019) such as attention (van Ede & Nobre, 2023) and decision-making (Pearson et al., 2014; Shadlen & Shohamy, 2016). A recent study demonstrated that human subjects can adeptly integrate information from working memory to drive decisions (van Ede & Nobre, 2024). Our results add to this growing body of evidence that working memory storage is a forward looking process – flexibly adapting based on expected future needs.

In closing, our findings suggest that what appears to be abstract deliberation may still be shaped by the structure of possible actions, and that working memory supports this flexibility not merely by retaining information, but by allowing deliberation to await its proper behavioral context.

## Data availability

Behavioral data and code to generate the figures will be uploaded to a publicly available repository on acceptance of the manuscript.

## Author Contributions

Conceptualization: SS, MNS

Formal Analysis: PS, SS

Funding Acquisition: MNS, SS

Investigation: PS

Methodology: SS, PS

Software: PS, SS, MNS

Supervision: SS, MNS

Writing - Original Draft Preparation: SS, PS

Writing - Review & Editing: SS, PS, MNS

## Acknowledgements

We thank Anne Loffler for help with online psychophysics testing that generated pilot data for this project.

